# Finding Nemo’s Genes: A chromosome-scale reference assembly of the genome of the orange clownfish *Amphiprion percula*

**DOI:** 10.1101/278267

**Authors:** Robert Lehmann, Damien J. Lightfoot, Celia Schunter, Craig T. Michell, Hajime Ohyanagi, Katsuhiko Mineta, Sylvain Foret, Michael L. Berumen, David J. Miller, Manuel Aranda, Takashi Gojobori, Philip L. Munday, Timothy Ravasi

**Author notes:** **Corresponding Author:** Timothy Ravasi, Division of Biological and Environmental Sciences & Engineering, King Abdullah University of Science and Technology, Thuwal, 23955-6900, Kingdom of Saudi Arabia.

## Abstract

The iconic orange clownfish, *Amphiprion percula*, is a model organism for studying the ecology and evolution of reef fishes, including patterns of population connectivity, sex change, social organization, habitat selection and adaptation to climate change. Notably, the orange clownfish is the only reef fish for which a complete larval dispersal kernel has been established and was the first fish species for which it was demonstrated that anti-predator responses of reef fishes could be impaired by ocean acidification. Despite its importance, molecular resources for this species remain scarce and until now it lacked a reference genome assembly. Here we present a *de novo* chromosome-scale assembly of the genome of the orange clownfish *Amphiprion percula*. We utilized single-molecule real-time sequencing technology from Pacific Biosciences to produce an initial polished assembly comprised of 1,414 contigs, with a contig N50 length of 1.86 Mb. Using Hi-C based chromatin contact maps, 98% of the genome assembly were placed into 24 chromosomes, resulting in a final assembly of 908.8 Mb in length with contig and scaffold N50s of 3.12 and 38.4 Mb, respectively. This makes it one of the most contiguous and complete fish genome assemblies currently available. The genome was annotated with 26,597 protein coding genes and contains 96% of the core set of conserved actinopterygian orthologs. The availability of this reference genome assembly as a community resource will further strengthen the role of the orange clownfish as a model species for research on the ecology and evolution of reef fishes.

## Introduction

The orange clownfish, *Amphiprion percula*, which was immortalized in the film “Finding Nemo”, is arguably the most recognized fish on Earth. It is also one of the most important species for studying the ecology and evolution of coral reef fishes. The orange clownfish is used as a model species to study patterns and processes of social organization (Buston, Bogdanowicz, Wong, & Harrison, 2007; Buston & Wong, 2014; Wong, Uppaluri, Medina, Seymour, & Buston, 2016), sex change (Buston, 2003), mutualism (Schmiege, D’Aloia, & Buston, 2017), habitat selection (Dixson et al., 2008; Elliott & Mariscal, 2001; Scott & Dixson, 2016), lifespan (Buston & García, 2007) and predator-prey interactions (Dixson, 2012; Manassa, Dixson, McCormick, & Chivers, 2013). It has been central to ground-breaking research into the scale of larval dispersal and population connectivity in marine fishes (Almany et al., 2017; Pinsky et al., 2017; Planes, Jones, & Thorrold, 2009; Salles et al., 2016) and how this influences the efficacy of marine protected areas (Berumen et al., 2012; Planes et al., 2009). It is also used to study the ecological effects of environmental disturbances in marine ecosystems (Hess, Wenger, Ainsworth, & Rummer, 2015; Wenger et al., 2014), including climate change (McLeod et al., 2013; Saenz-Agudelo, Jones, Thorrold, & Planes, 2011) and ocean acidification (Dixson, Munday, & Jones, 2010; Jarrold, Humphrey, McCormick, & Munday, 2017; Munday et al., 2009; Simpson et al., 2011). Perhaps more than any other species, the orange clownfish has become a mainstay of research into the chemical, molecular, behavioral, population, conservation and climate-change ecology of marine fishes.

The orange clownfish is one of 30 species of anemonefishes belonging to the subfamily Amphiprioninae within the family Pomacentridae (damselfishes). The two clownfishes, *A. percula* (orange clownfish or clown anemonefish) and *A. ocellaris* (false clownfish or western clown anemonefish) form a separate clade, alongside *Premnas biaculeatus*, within the Amphiprioninae (J. Li, Chen, Kang, & Liu, 2015; Litsios, Pearman, Lanterbecq, Tolou, & Salamin, 2014; Litsios & Salamin, 2014). The two species of clownfish are easily distinguished from other anemonefishes by their bright orange body coloration and three vertical white bars. The orange clownfish and the false clownfish have similar body coloration, but largely distinct allopatric geographical distributions (Litsios & Salamin, 2014). The orange clownfish occurs in northern Australia, including the Great Barrier Reef (GBR), and in Papua New Guinea, Solomon Islands and Vanuatu, while the false clownfish occurs in the Indo-Malaysian region, from the Ryukyu Islands of Japan, throughout south-east Asia and south to north-western Australia (but not the GBR).

Like all anemonefishes, the orange clownfish has a mutualistic relationship with sea-anemones. Wild adults and juveniles live exclusively in association with a sea anemone, where they gain shelter from predators and benefit from food captured by the anemone (Fautin, 1991; Fautin & Allen, 1997; Mebs, 2009). In return, the sea-anemone benefits by gaining protection from predators (Fautin & Allen, 1997; Holbrook & Schmitt, 2005), from supplemental nutrition from the clownfish’s waste (Holbrook & Schmitt, 2005) and from increased gas exchange as a result of increased water flow provided by clownfish movement and activity (Herbert, Bröhl, Springer, & Kunzmann, 2017; Szczebak, Henry, Al-Horani, & Chadwick, 2013). The orange clownfish associates with two species of anemone, *Stichodactyla gigantea* and *Heteractis magnifica* (Fautin & Allen, 1997). Clownfish social groups typically consist of an adult breeding pair and a variable number of smaller, size-ranked juveniles that queue for breeding rights (Buston, 2003). The breeding female is larger than the male. If the female disappears, the male changes sex to female and the largest non-breeder matures into a breeding male. The breeding pair lays clutches of demersal eggs in close proximity to their host anemone. Eggs hatch after 7-8 days and the larvae disperse into the open ocean for a period of 11-12 days, at which time they return to the reef and settle to an anemone.

The close association of clownfish and other anemonefishes with sea anemones makes them excellent species for studying aspects of marine mutualisms and habitat selection. The easily identified and delineated habitat they occupy, along with the ease with which the fish can be observed in nature, makes them ideal candidates for behavioral and population ecology. The unique capacity to collect juveniles immediately after they have settled to the reef from their pelagic larval phase also makes them ideally suited to testing long-standing questions about larval dispersal and population connectivity in reef fish populations. Using molecular techniques to assign parentage between newly settled juveniles and adult anemonefishes, recent studies have been able to describe for the first time the spatial scales of dispersal in reef fish and its temporal consistency (Almany et al., 2017). The ability to map the connectivity of clownfish populations in space and time has also opened the door to addressing challenging questions about selection, fitness and adaptation in natural populations of marine fishes (Pinsky et al., 2017; Salles et al., 2016). Finally, the orange clownfish is one of the relatively few coral reef fishes that can easily be reared in captivity (Wittenrich, Turingan, & Creswell, 2007). Consequently, it has unrivalled potential for experimental manipulation to test ecological and evolutionary questions in marine ecology (Dixson et al., 2014; Manassa et al., 2013), including the impacts of climate change and ocean acidification (Nilsson et al., 2012). Increasingly, genome-wide methods are being used to test ecological and evolutionary questions and this is particularly true for coral reef species in the wake of anthropomorphic climate change and its effects on these sensitive ecosystems (Stillman & Armstrong, 2015)

To date, genome assemblies of two anemonefish, *A. frenatus* (Marcionetti, Rossier, Bertrand, Litsios, & Salamin, 2018) and *A. ocellaris* (Marcionetti et al., 2018), have been published. Both of these were based on short-read Illumina technology with genome scaffolding provided by shallow coverage of PacBio (Marcionetti et al., 2018) or Oxford Nanopore (Tan et al., 2018) long reads. While the use of long reads to scaffold Illumina-based assemblies improves contiguity, both genome assemblies are highly fragmented with respective contig and scaffold N50s of 14.9 and 244.5 kb for *A. frenatus* and 323.6 and 401.7 kb for *A. ocellaris*. Here we present a chromosome-scale genome assembly of the orange clownfish, which was assembled using a primary PacBio long read strategy, followed by scaffolding with Hi-C-based chromatin contact maps. The resulting final assembly is highly contiguous with contig and scaffold N50 values of 3.12 and 38.4 Mb, respectively. This assembly will be a valuable resource for the research community and will further establish the orange clownfish as a model organism for genetic and genomic studies into ecological, evolutionary and environmental aspects of reef fishes. To facilitate the use of this resource, we have developed an integrated database, the Nemo Genome DB (http://nemogenome.org/), which allows for the interrogation and mining of genomic and transcriptomic data described here.

## Materials and Methods

### Specimen collection and DNA extraction

Adult orange clownfish breeding pairs were collected on the northern GBR in Australia. Fish were bred at the Experimental Aquarium Facility of James Cook University (JCU) and one individual offspring was sacrificed at the age of 8 months. The whole brain was excised, snap frozen and kept at −80°C until processing. High molecular weight DNA was extracted from whole brain tissue using the Qiagen Genomic-tip 100/G extraction kit. The tissue was first homogenized in lysis buffer G2 supplemented with 200 μg/mL RNase A using sterile beads for 30 sec. After homogenization, proteinase K was added and the homogenate was incubated at 50°C overnight. DNA extraction was then performed according to the manufacturer’s protocol with a final elution volume of 200 μl. DNA fragment size and quality was assessed using pulsed-field gel electrophoresis. This study was completed under JCU animal ethics permits A1961 and A2255.

### PacBio library preparation and sequencing

For Pacific Biosciences (PacBio) long read sequencing, the extracted orange clownfish DNA was first sheared using a g-TUBE (Covaris, MA, USA) (target size of 20 kb) and then converted into SMRTbell template libraries according to the manufacturer’s protocol (Pacific Biosciences, CA, USA). Size selection was performed using BluePippin (Sage Science, MA, USA) to generate two libraries with a minimum size of 10 and 15 kb, respectively. Sequencing was performed using P6-C4 chemistry on the PacBio RS II instrument at the King Abdullah University of Science and Technology (KAUST) Bioscience Core Laboratory (BCL) with 360 mins movies. A total of 113 SMRT cells were sequenced.

### Genome assembly

The genome sequence was assembled from the unprocessed PacBio reads (Table S1) using the hierarchical diploid aware PacBio assembler FALCON v0.4.0 (Chin et al., 2016). To obtain the optimal assembly, different parameters were tested (Table S2) to generate 12 candidate assemblies. The contiguity of these assemblies was assessed with QUAST v3.2 (Gurevich, Saveliev, Vyahhi, & Tesler, 2013), while assembly completeness was determined with BUSCO v2.0 (Simão, Waterhouse, Ioannidis, Kriventseva, & Zdobnov, 2015). Assembly “A7” exhibits the highest contiguity and single copy orthologous gene completeness and was selected for further improvement. The FALCON_Unzip algorithm was then applied to the initial A7 assembly obtain a haplotype-resolved, phased assembly, termed “A7-phased”. Contigs less than 20 kb in length were removed from the assembly. This phased assembly was polished with Quiver to achieve final consensus sequence accuracies comparable to Sanger sequencing (Chin et al., 2013) using default settings, which produced the “A7-phased-polished” assembly.

### Genome assembly scaffolding with chromatin contact maps

The flash-frozen brain tissue was sent to Phase Genomics (Seattle, WA, USA) for the construction chromatin contact maps. Tissue fixation, chromatin isolation, library preparation and 80-bp paired end sequencing were performed by Phase Genomics. The sequencing reads were aligned to the A7-phased-polished version of the assembly with BWA (H. Li & Durbin, 2010) and uniquely mapping read pairs were retained. Contigs from the A7-phased-polished assembly were clustered, ordered and then oriented using Proximo (Bickhart et al., 2017; Burton et al., 2013), with settings as previously described (Peichel, Sullivan, Liachko, & White, 2017). Briefly, contigs were clustered into chromosomal groups using a hierarchical clustering algorithm based on the number of read pairs linking scaffolds, with the final number of groups specified as the number of the haploid chromosomes. The haploid chromosome number was set as 24, which is consistent with the observed haploid chromosome number of the Amphiprioninae, as published for *A. ocellaris* (Arai, Inoue, & Ida, 1976), *A. frenatus*, (Molina & Galetti, 2004; Takai & Kosuga, 2007), *A. clarkii* (Arai & Inoue, 1976; Takai & Kosuga, 2007), *A. perideraion* (Supiwong et al., 2015) and *A. polymnus* (Tanomtong et al., 2012). After clustering into chromosomal groups, the scaffolds were ordered based on Hi-C link densities and then oriented with respect to the adjacent scaffolds using a weighted directed acyclic graph of all possible orientations based on the exact locations of the Hi-C links between scaffolds. Gaps between contigs were represented with 100 Ns and the proximity-guided assembly was named “A7-PGA”. Gaps in the scaffolded assembly were subsequently closed using PBJelly from PBSuite v15.8.24 (English et al., 2012) with the entire PacBio read dataset and Blasr (Chaisson & Tesler, 2012) (parameters: --minMatch 8 --minPctIdentity 70 --bestn 1 --nCandidates 20 --maxScore −500 --nproc 32 --noSplitSubreads), to give rise to the final version of the assembly, “Nemo v1”.

### Genome assembly validation

Genomic DNA was extracted from a second individual and Illumina sequencing libraries were prepared using the NEBNext Ultra II DNA library prep kit for Illumina following the manufacturer’s protocol. Three cycles of PCR were used to enrich the library. The sequencing libraries were sequenced on two lanes of a HiSeq 2500 at the KAUST BCL. A total of 1,199,533,204 paired reads were generated, covering approximately 181 Gb. The 151-bp paired end reads were processed with Trimmomatic v0.33 to remove adapter sequences and low-quality stretches of nucleotides (parameters: 2:30:10 LEADING:20 TRAILING:20 SLIDINGWINDOW:4:20 MINLEN:75) (Bolger, Lohse, & Usadel, 2014).

The genome assembly size was validated by comparison to a k-mer based estimate of genome size. The first half of the paired-end reads of one sequencing lane (∼ 25 Gb of data) was used for the k-mer estimate of genome size. Firstly, KmerGenie (Chikhi & Medvedev, 2014) was used to determine the optimal k-value for a k-mer based estimation. Following that, Jellyfish v2.2.6 (Marçais & Kingsford, 2011) was used with k=71 to obtain the frequency distribution of all k-mers with this length. The resulting distribution was analyzed with Genomescope (Vurture et al., 2017) to estimate genome size, repeat content and the level of heterozygosity. To further validate the assembly, we determined the proportion of trimmed Illumina short reads that mapped to the Nemo v1 assembly with BWA v0.7.10 (H. Li & Durbin, 2010) and SAMtools v1.1 (H. Li et al., 2009). Additionally, the completeness of the genome assembly annotation as determined by the conservation of a core set of genes was measured using BUSCO with default parameters.

### Repeat annotation

A species-specific *de novo* repeat library was assembled by combining the results of three distinct repeat annotation methods. Firstly, RepeatModeler v1.08 (Smit & Hubley, 2008) was used to build an initial repeat library. Secondly, we used LtrHarvest (Ellinghaus, Kurtz, & Willhoeft, 2008) and LTRdigest (Steinbiss, Willhoeft, Gremme, & Kurtz, 2009), both accessed *via* genometools 1.5.6 (Gremme, Steinbiss, & Kurtz, 2013), with the following parameters: -seed 76 -xdrop 7 -mat 2 -mis −2 -ins −3 -del −3 -mintsd 4 -maxtsd 20 -minlenltr 100 -maxlenltr 6000 -maxdistltr 25000 -mindistltr 1500 -similar 90. The resulting hits were filtered with LTRdigest, accepting only sequences featuring a hit to one of the hidden markov models in the GyDB 2.0 database. Thirdly, TransposonPSI v08222010 (Haas, 2018) was used to detect sequences with similarities to known families of transposon open reading frames. To remove duplicated sequences in the combined result from all three methods a clustering with USEARCH (Edgar, 2010) was performed requiring at least 90% sequence identity, and only cluster representatives were retained. The resulting representative sequences were classified by RepeatClassifier (part of RepeatModeler), Censor v4.2.29 (Jurka, Klonowski, Dagman, & Pelton, 1996) and Dfam v2.0 (Wheeler et al., 2012), and were then blasted against the Uniprot/Swissprot database (release 2017_12) to obtain a unified classification. Furthermore, these three classification methods and the blast result was used to filter out spurious matches to protein-coding sequence. Specifically, putative repeat sequences were only retained when at least one classification method recognized the sequence as a repeat and the best match in Swissprot/Uniprot was not a protein-coding gene (default blastx settings). Furthermore, sequences were retained if two of the three identification methods classified the sequence as repeat, but the best blast hit was not a transposable element. This *de novo* library was combined with the thoroughly curated zebrafish repeat library provided by Repbase v22.05 (Bao, Kojima, & Kohany, 2015) and this combined library was employed for repeat masking in the Nemo v1 assembly using RepeatMasker (Smit, Hubley, & Green, 2010).

### RNA extraction, library construction, sequencing and read processing

Tissues for RNA extraction were dissected from one eight-month old orange clownfish individual. RNA was extracted from skin, eye, muscle, gill, liver, kidney, gallbladder, stomach and fin tissues using the Qiagen AllPrep kit following manufacturer’s instructions. Sequencing libraries were prepared using the TruSeq Stranded mRNA Library Preparation kit and 150 bp paired-end sequencing was performed on one lane of an Illumina HiSeq 4000 machine in the KAUST BCL. The RNA-seq reads were trimmed with Trimmomatic v0.33 (Bolger et al., 2014) (parameters: 2:30:10 LEADING:3 TRAILING:3 SLIDINGWINDOW:4:15 MINLEN:40) and contamination was removed with Kraken (Wood & Salzberg, 2014) by retaining only unclassified reads.

### Genome assembly annotation

After mapping the RNA-seq data with STAR v2.5.2b (Dobin et al., 2013) to the final assembly, an *ab-initio* annotation with BRAKER1 v1.9 (Hoff, Lange, Lomsadze, Borodovsky, & Stanke, 2016) was performed. This initial annotation identified 49,881 genes. This annotation was then integrated with external evidence using the MAKER2 v2.31.8 (Holt & Yandell, 2011) gene annotation pipeline. First, the transcriptome of the orange clownfish was provided to MAKER2 as EST evidence in two forms, a *de novo* assembly of the preprocessed RNA-seq reads obtained with Trinity v2.4.0 (Grabherr et al., 2011), and a genome-guided assembly performed with the Hisat2 v2.1.0/Stringtie v1.3.3b workflow (Pertea, Kim, Pertea, Leek, & Salzberg, 2016). Second, we combined the proteomes of zebrafish (GCF_000002035.6_GRCz11), Nile tilapia (GCF_001858045.1_ASM185804v2) and bicolor damselfish (*Stegastes partitus*) (GCA_000690725.1), together with the Uniprot/Swissprot database (release 2017_12: 554,515 sequences) and the successfully detected BUSCO genes to generate a reference protein set for homology based gene prediction. In the initial MAKER2 run, the annotation edit distances (AED) were calculated for the BRAKER1-obtained annotation, and only gene annotations with an AED of less than 0.1 and a corresponding protein length of greater than 50 amino acids were retained for subsequent training of the gene prediction program SNAP v2013.11.29 (Korf, 2004). Similarly, the AUGUSTUS v3.2.3 (Stanke et al., 2006) gene prediction program was trained on 1,850 gene annotations that possessed: an AED score of less than 0.01; an initial start codon, a terminal stop codon and no in-frame stop codons; more than one exon; and no introns greater than 10 kb. The hidden markov gene model of GeneMark v4.32 (Ter-Hovhannisyan, Lomsadze, Chernoff, & Borodovsky, 2008) was trained by BRAKER1. The final annotation was then obtained in the second run of MAKER2 with the trained models for SNAP, GeneMark, and AUGUSTUS. InterProScan 5 was then used to obtain the Pfam protein domain annotations for all genes. The standard gene builds were then generated. The output was filtered to include all annotated genes with evidence (AED less than 1) or with a Pfam protein domain, as recommended (Campbell, Holt, Moore, & Yandell, 2014).

### Functional annotation

The protein sequences produced from the genome assembly annotation were aligned to the UniProtKB/Swiss-Prot database (release 2017_12) with blastp v2.2.29 (parameters: -outfmt 5-evalue 1e-3 -word_size 3 -show_gis -num_alignments 20 -max_hsps 20) and protein signatures were annotated with InterProScan 5. The results were then integrated with Blast2GO v4.1.9 (Gotz et al., 2008).

### Genome assembly comparisons

For genome assembly comparisons, we compared the Nemo v1 genome assembly to the 26 previously reported fish chromosome-scale genome assemblies (Table S3). Comparisons were made for genome assembly contiguity and completeness. Contig N50 values are reported for the scaffold-scale versions of each assembly and are taken from the indicated publication (Table S3), database description (Table S3) or were generated with the Perl assemblathon_stats_2.pl script (Bradnam et al., 2013). Genome assembly completeness was assessed by determining the proportion of the genome size that is contained within the chromosome content of each assembly. It should be noted that this comparison is relative to the estimated genome size and not the published assembly size. The estimated genome size was taken as either the published estimated genome size in the relevant paper (Table S3), or from the Animal Genome Size Database (Gregory, 2018). Where possible, k-mer derived or flow cytometry-based estimates of genome size were used. Before calculation, we remove stretches of Ns from the genome assemblies as these are used to arbitrarily space scaffolds and do not contain actual genome information. However, this step was not possible for the Asian arowana, southern platyfish, yellowtail or croaker genomes as the chromosome-scale assemblies have not been made publicly available. Genome assembly completeness was determined with BUSCO (Simão et al., 2015) using the Actinopterygii set of 4,584 genes and the AUGUSTUS zebrafish gene model provided with the software.

### Gene homology

To investigate the gene space of the orange clownfish genome assembly, we used OrthoFinder v1.1.4 (Emms & Kelly, 2015) to identify orthologous gene relationships between the orange clownfish and four related fish species. The following four fish species were utilized in addition to the orange clownfish: Asian seabass GCF_001640805.1_ASM164080v1 (45,223 sequences), Nile tilapia GCF_001858045.1_ASM185804v2 (58,087 sequences), southern platyfish GCF_000241075.1_*Xiphophorus_maculatus*-4.4.2 (23,478 sequences), and zebrafish GCF_000002035.6_GRCz11 (52,829 sequences). The longest isoform of each gene was utilized in the analysis, which corresponded to 25,050, 28,497, 23,043, and 32,420 sequences, respectively. 26,597 sequences were used for the orange clownfish. These protein sequences were reciprocally blasted against each other and clusters of orthologous genes were then defined using OrthoFinder with default parameters. As part of OrthoFinder, the concatenated sequences of single-copy orthologs present in all species were then used to construct a phylogenetic tree, which was rooted using STRIDE (Emms & Kelly, 2017).

### Database system architecture and software

The Nemo Genome DB database (http://nemogenome.org) was implemented on a UNIX server with CentOS version 7, Apache web server and MySQL Database server. JBrowse (Buels et al., 2016) was employed to visualize the genome assembly and genomic features graphically and interactively. JavaScript was adopted to implement client-side rich applications. The JavaScript library, jQuery (http://jquery.com) was employed. Other conventional utilities for UNIX computing were appropriately installed on the server if necessary. All of the Nemo Genome DB resources are stored on the server and are available through HTTP access.

## Results and Discussion

### Sequencing and assembly of the orange clownfish genome

Genomic DNA of an individual orange clownfish (Fig. 1A) was sequenced with the PacBio RS II platform to generate 1,995,360 long reads, yielding 113.8 Gb, which corresponds to a 121-fold coverage of the genome (Table S1). After filtering with the read pre-assembly step of the Falcon assembler, 5,764,748 reads, covering 54.3 Gb and representing a 58-fold coverage of the genome, were available for assembly.

**Fig. 1.**
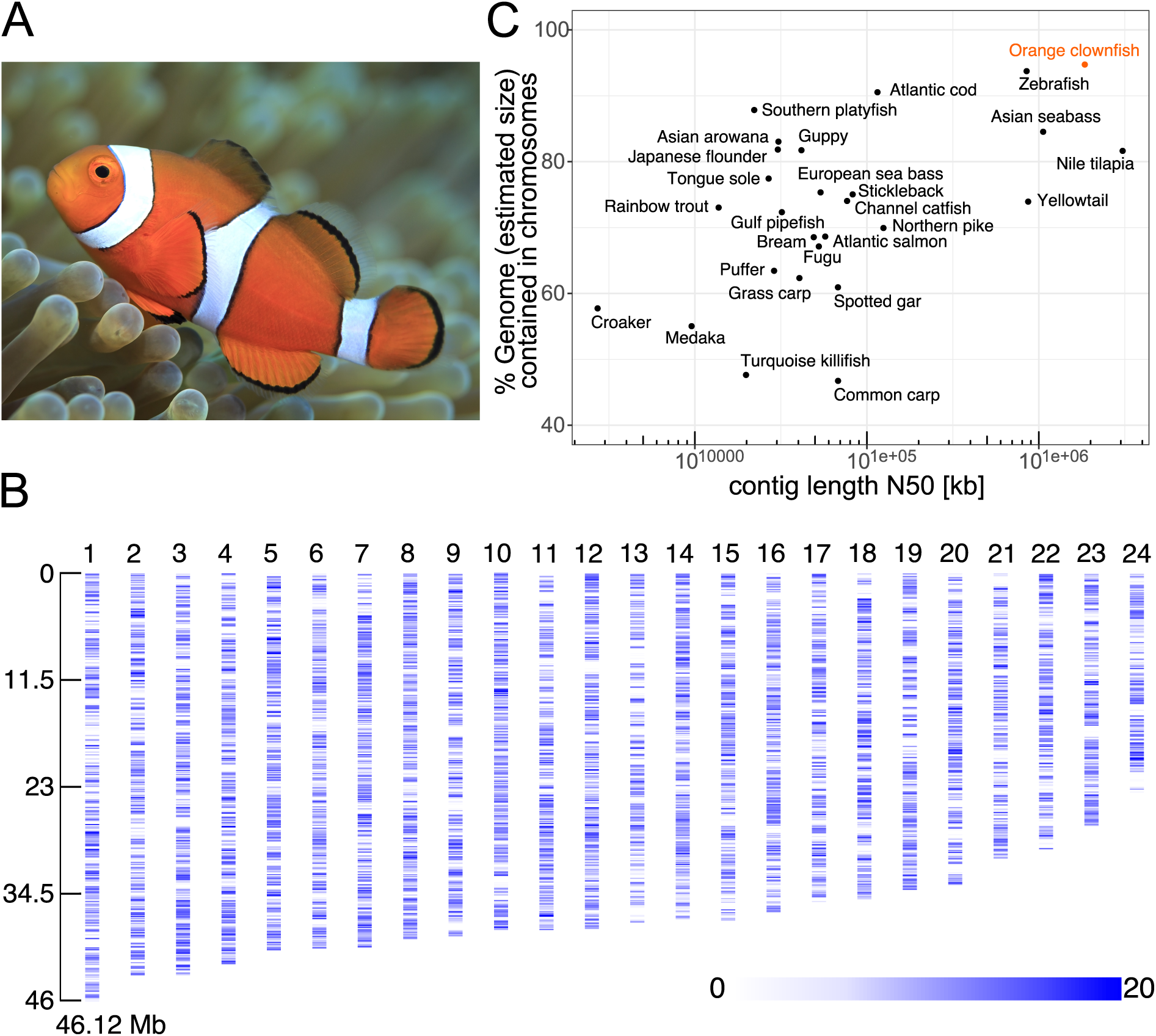
**(A)** The iconic orange clownfish (*A. percula*). **(B)** Gene density on the 24 chromosomes, plotted in 100 kb windows. Chromosomes are ordered by size, as indicated on the left axis in Mb. **(C)** Contiguity (x-axis) and genome assembly completeness (y-axis) of the orange clownfish, and the 26 previously published, chromosome-scale fish genome assemblies. Details and statistics of the 27 assemblies are presented in Table S3.

To optimize the assembly parameters, we performed 12 trial assemblies using a range of parameters for different stages of the Falcon assembler (Table S2). The assembly quality was assessed by considering assembly contiguity (contig N50 and L50), total assembly size, and also gene completeness (BUSCO) (Table 1). Assembly A7 exhibited the highest contig N50 (1.80 Mb), lowest contig L50 (138 contigs), lowest number of missing BUSCO genes (132) and is only slightly surpassed in the longest contig metric (15.8 Mb) by the highly similar assemblies A8 and A9 (16.5 Mb) (Table 1).

**Table 1.** Contig statistics for the preliminary candidate-assemblies

| Assembly | Length<br>(Mb) | Number | N50<br>(Mb) | L50 | Longest<br>(Mb) | Missing genes<br>(number (%)) |
| --- | --- | --- | --- | --- | --- | --- |
| A1 | 950.4 | 4,874 | 1.024 | 254 | 9.59 | 148 (3.23) |
| A2 | 945.4 | 4,374 | 1.040 | 251 | 6.67 | 156 (3.30) |
| A3 | 926.5 | 3,629 | 1.070 | 236 | 7.21 | 140 (3.05) |
| A4 | 921.8 | 2,829 | 1.380 | 184 | 8.16 | 134 (2.92) |
| A5 | 883.9 | 1,017 | 1.469 | 167 | 10.24 | 146 (3.18) |
| A6 | 902.2 | 2,204 | 1.401 | 174 | 12.38 | 134 (2.92) |
| A7 | 920.7 | 2,473 | 1.801 | 138 | 15.84 | 132 (2.88) |
| A8 | 924.6 | 2,629 | 1.742 | 143 | 16.51 | 139 (3.03) |
| A9 | 924.9 | 2,638 | 1.742 | 143 | 16.51 | 140 (3.05) |
| A10 | 917.1 | 2,368 | 1.648 | 140 | 10.21 | 146 (3.18) |
| A11 | 899.9 | 2,049 | 1.571 | 160 | 9.07 | 151 (3.29) |
| A12 | 908.8 | 2,086 | 1.602 | 142 | 10.21 | 143 (3.12) |

Genome assemblies represent a mixture of the two possible haplotypes of a diploid individual at each locus. This collapsing of haplotypes may result in a loss of important sequence information. However, diploid-aware assembly algorithms such as the Falcon_Unzip assembler are designed to detect single nucleotide polymorphisms (SNPs) as well as structural variations and to use this information to phase (“unzip”) heterozygous regions into distinct haplotypes (Chin et al., 2016). This procedure results in a primary assembly and a set of associated haplotype contigs (haplotigs) capturing the divergent sequences. Having established the parameter set that gave the best assembly metrics with Falcon, we used Falcon_Unzip to produce a phased assembly (“A7-phased”) of the orange clownfish (Table 2). The phased assembly was 905.0 Mb in length with a contig N50 of 1.85 Mb. As has been seen in previous genome assembly projects (Chin et al., 2016), Falcon_Unzip produced a smaller assembly with fewer contigs than the assembly produced by Falcon (Table 2). The phased primary assembly was then polished with Quiver, which yielded an assembly (“A7-phased-polished”) with 1,414 contigs spanning 903.6 Mb with an N50 of 1.86 Mb (Table 2). This polishing step closed 91 gaps in the assembly and improved the N50 by approximately 14.3 kb. After polishing of the “unzipped” A7-phased-polished assembly, 9,971 secondary contigs were resolved, covering 340.1 Mb of the genome assembly. The contig N50 of these secondary contigs was 38.2 kb, with over 99% of them being longer than 10 kb in size. Relative to the 903.6 Mb A7-phased-polished primary contig assembly, the secondary contigs covered 38% of the assembly size. To the best of our knowledge, this is the first published fish genome assembly that has been resolved to the haplotype level with Falcon_Unzip.

**Table 2.**
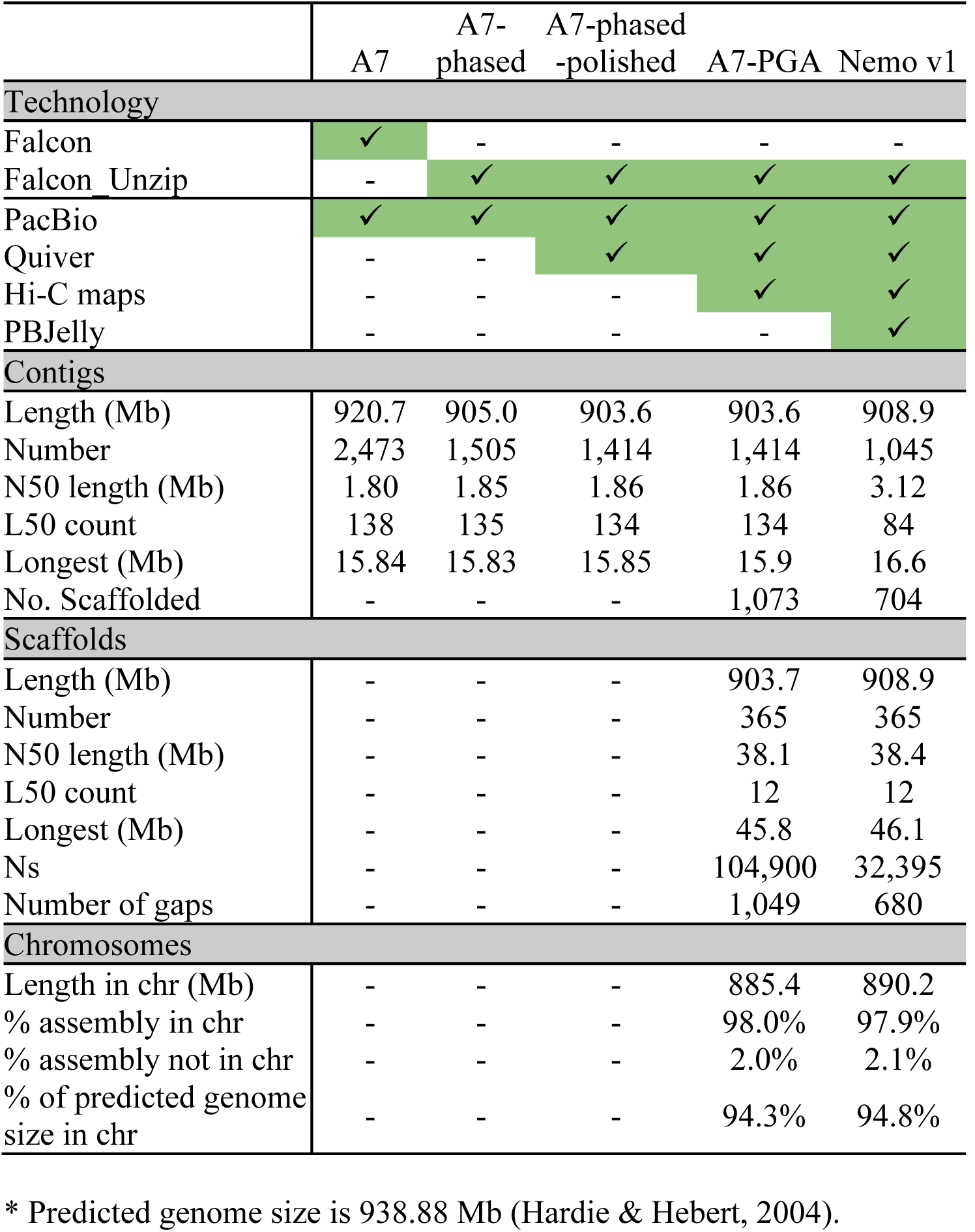
Assembly statistics of the orange clownfish genome assemblies

### Scaffolding of the orange clownfish genome assembly into chromosomes

To build a chromosome-scale reference genome assembly of the orange clownfish, chromatin contact maps were generated by Phase Genomics (Fig. S1). Scaffolding was performed by the Proximo algorithm (Bickhart et al., 2017; Burton et al., 2013) on the A7-phased-polished assembly using 231 million Hi-C-based paired-end reads to produce the proximity guided assembly “A7-PGA” (Table 2). The contig clustering allowed the placement of 1,073 contigs into 24 scaffolds (chromosomes) with lengths ranging from 23.4 to 45.8 Mb (Tables 2 and 3). While only 76% of the contigs were assembled into chromosome clusters, this corresponds to 98% (885.4 Mb) of total assembly length and represents 95% of the estimated genome size of 938.9 Mb (Tables 2 and 3). This step substantially improved the overall assembly contiguity, raising the N50 20-fold from 1.86 to 38.1 Mb.

**Table 3.**
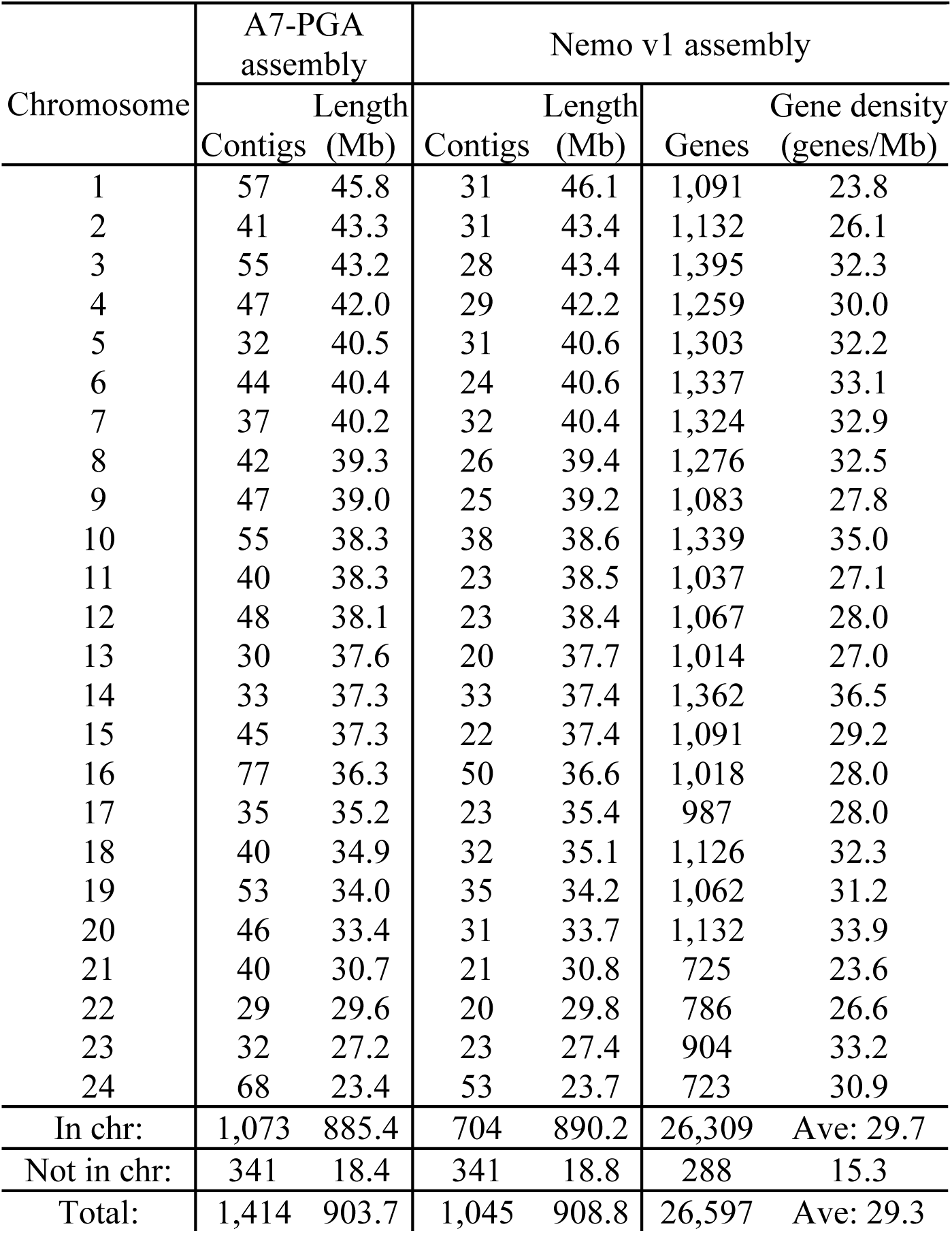
Chromosome metrics before and after polishing of the final assembly

A quality score for the order and orientation of contigs within the A7-PGA assembly was determined. This metric is based on the differential log-likelihood of the contig orientation having produced the observed log-likelihood, relative to its neighbors (Burton et al., 2013). The orientation of a contig was deemed to be of high quality if its placement and orientation, relative to neighbors, was 100 times more likely than alternatives (Burton et al., 2013). In A7-PGA, the placements of 524 (37%) of the scaffolds were deemed to be of high quality, accounting for 775.5 Mb (87%) of the scaffolded chromosomes, indicating the robustness of the assembly.

A final polishing step was performed with PBJelly to generate the final Nemo v1 assembly. This polishing step closed 369 gaps, thereby improving the contig N50 by 68% and increasing the total assembly length by 5.21 Mb (Tables 2 and 3). The length of each chromosome was increased, with a range of 23.7 to 46.1 Mb (Fig. 1B). Gaps were closed in each chromosome except for chromosome 14, leaving an average of only 28 gaps per chromosome (Table 3). The final assembly is 908.9 Mb in size and has contig and scaffold N50s of 3.12 and 38.4 Mb, respectively. The assembly is highly contiguous as can be observed by the fact that 50% of the genome length is contained within the largest 84 contigs. 890.2 Mb (98%) of the genome assembly size was scaffolded into 24 chromosomes, with only 18.8 Mb of the assembly failing to be grouped. The 18.8 Mb of unscaffolded assembly is comprised of 341 contigs with a contig N50 of only 57.8 kb.

### Validation of the orange clownfish genome assembly size

The final assembly size of 908.9 Mb is consistent with the results of a Feulgen image analysis densitometry-based study, which determined a C-value of 0.96 pg and thus a genome size of 938.9 Mb for the orange clownfish (Hardie & Hebert, 2004). Furthermore, our assembly size is in keeping with estimates of genome size for other fish of the *Amphiprion* genus, which range from 792 to 1193 Mb (Gregory, 2018). We additionally validated the observed assembly size by using a k-mer based approach. Specifically, the k-mer coverage and frequency distribution were plotted and fitted with a four-component statistical model with GenomeScope (Fig. S2A). This allowed us to generate an estimate of genome size as well as the repeat content and level of heterozygosity. However, varying the k-value from the recommended value of 21 up to 27 yielded a corresponding increase of the estimated genome size. We therefore used KmerGenie to determine the optimal k-mer length of 71 to capture the available sequence information. The utilization of small k-values might partially explain the reported tendency of GenomeScope to underestimate the genome size (Vurture et al., 2017). The final estimate of the haploid genome length by k-mer analysis was 906.6 Mb, with 732.8 Mb (80%) of unique sequence and a repeat content of 173.8 Mb (19%). Furthermore, the estimated heterozygosity level of 0.12% is low considering that an F1 offspring of wild caught fish was sequenced (Fig. S2B). While the short-read k-mer based genome size estimate of 906.6 Mb matches the final assembly size of 908.9 Mb very well, the C-value derived genome size estimate is slightly larger (938.9 Mb). As an additional validation of the accuracy of the genome assembly, we mapped the trimmed Illumina short reads to the Nemo v1 assembly and observed that 95% of the reads mapped to the assembly and that 84% of the reads were properly paired.

Based on the C-value derived genome size estimate, there is approximately 29.9 Mb (3.3%) of sequence length absent from our genome assembly. It seems likely that our assembly is nearly complete for the euchromatic regions of the genome given our assessment of genome size and gene content completeness. However, genomic regions such as the proximal and distal boundaries of euchromatic regions contain heterochromatic and telomeric repeats, respectively, are refractory to currently available sequencing techniques and are typically absent from genome assemblies (Bickhart et al., 2017; Hoskins et al., 2007).

### Chromosome-scale fish genome assembly comparisons

To date, chromosome-scale genome assemblies have been released for 26 other fish species (Table S3). Here, we present the first chromosome-scale assembly of a tropical coral reef fish, the orange clownfish. As a measure of genome assembly quality, we assessed the contiguity and completeness of these 27 chromosome-scale genome assemblies. We investigated genome contiguity with the contig N50 metric and characterized genome completeness for each genome assembly by calculating the proportion of the estimated genome size that was assigned to chromosomes. As shown in Fig. 1C, the orange clownfish genome assembly is highly contiguous, with a scaffold-scale contig N50 of 1.86 Mb, which is only surpassed by the contig N50 of the Nile tilapia genome assembly. Interestingly, even though different assembler algorithms were utilized, the three genome assemblies based primarily on long read PacBio technology were the most contiguous, with only Nile tilapia (3.09 Mb, Canu), orange clownfish (1.86 Mb, Falcon) and Asian seabass (1.19 Mb, HGAP) genome assemblies yielding contig N50s in excess of 1 Mb.

While the use of long read sequencing technologies facilitates the production of highly contiguous genome assemblies, scaffold sizes are still much shorter than the length of the underlying chromosomes. The use of further scaffolding technologies such as genetic linkage maps, scaffolding based on synteny with genome assemblies from related organisms, as well as *in vitro* and *in vivo* Hi-C based methods has allowed for the production of assemblies with chromosome-sized scaffolds. Here, the use of Hi-C based chromatin contact maps allowed for the placement of 98% of the Nemo v1 assembly length (890.2 of 908.9 Mb) into chromosomes, yielding a final assembly with a scaffold N50 of 38.4 Mb. This corresponds to 95% of the estimated genome size (938.9 Mb), which suggests that the Nemo v1 assembly is one of the most complete fish genome assemblies published to date (Fig. 1C). Only the zebrafish (94%) and Atlantic cod (91%) genome assemblies had a comparably high proportion of their estimated genome sizes scaffolded into chromosome-length scaffolds (Fig. 1C). It is likely that the use of both PacBio long reads and Hi-C based chromatin contact maps contributed to the very high proportion of the orange clownfish genome that we were able to both sequence and assemble into chromosomes.

While assembly contiguity is important, genome completeness with respect to gene content is also vital for producing a genome assembly that will be utilized by the research community. We evaluated the completeness of the 27 chromosome-scale assemblies with BUSCO and the Actinopterygii lineage, which encompasses 4,584 highly-conserved genes. When ranked by the total of complete (single copy and duplicate) genes, the orange clownfish assembly is the second most complete, with 4,456 (97.2%) of the orthologs identified (Fig. 2). The top ranked assembly, Nile tilapia, contains only 9 more of the core set of orthologs such that it contains 4,465 of the orthologs (97.4%). While the assemblies based on PacBio long read technology are again amongst the most complete, it should also be noted that most of the assemblies analyzed showed a very high level of completeness.

**Fig. 2.**
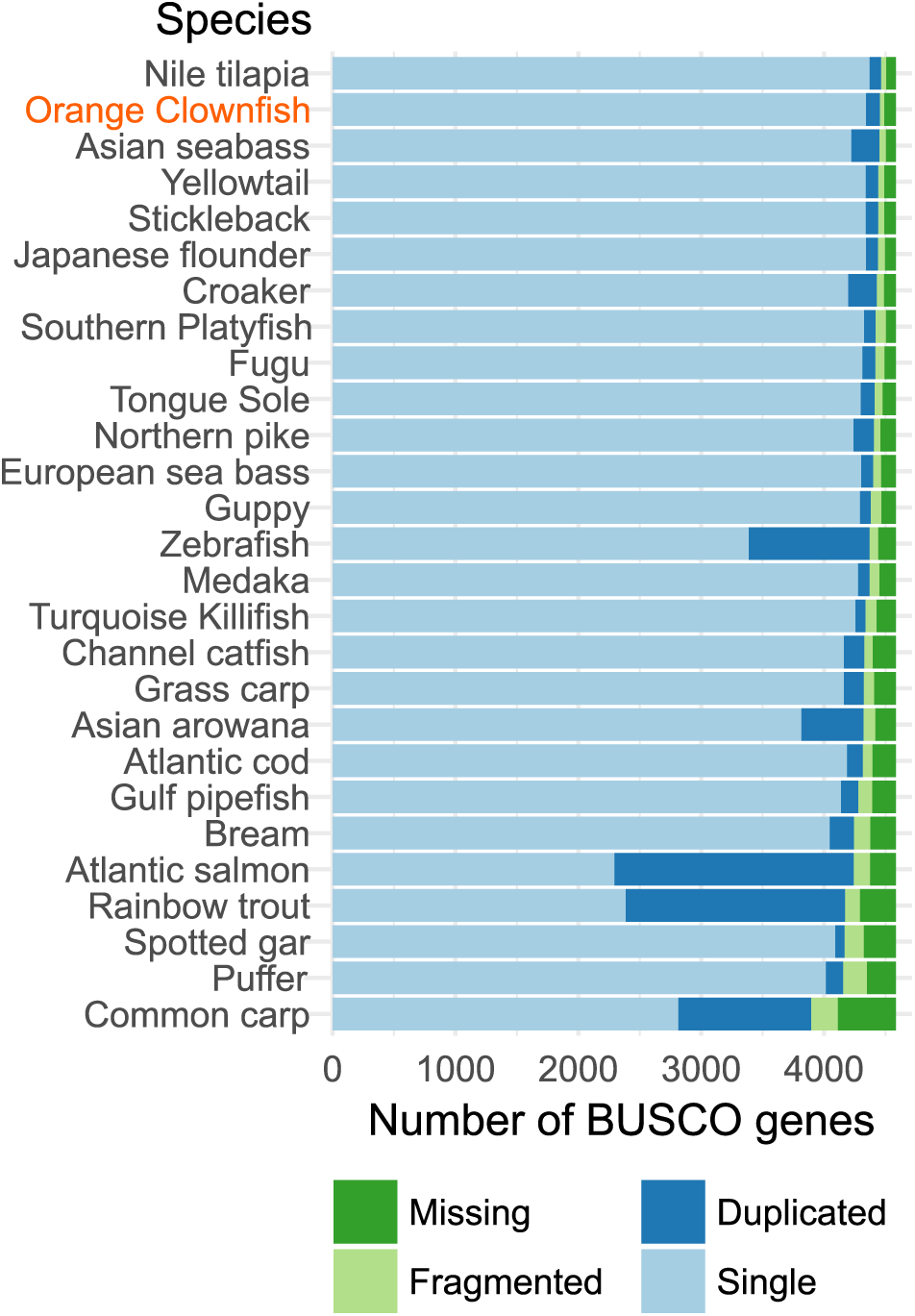
Genome assembly completeness of all published chromosome-scale fish genome assemblies, as measured by the proportion of the BUSCO set of core genes detected in each assembly. Genome assemblies on the y-axis are sorted by the sum of single copy and duplicated BUSCO genes.

### Genome annotation

To annotate repetitive sequences and transposable elements, we constructed an orange clownfish-specific library by combining the results of Repeatmodeler, LTRharvest and TransposonPSI. Duplicate sequences were removed and false positives were identified using three classification protocols (Censor, Dfam, RepeatClassifier) as well as comparisons to Uniprot/Swissprot databases. After these filtering steps, we identified 21,644 repetitive sequences. These sequences, in combination with the zebrafish library of RepBase, were then used for genome masking with RepeatMasker. This lead to a total of 28% of the assembly being identified as repetitive (Fig. 3A and Table S4). It was observed that there is a general trend for increased repeat density towards the ends of chromosome arms (Fig. 3B and S2). The total fraction of repetitive genomic sequence is in good agreement with other related fish species (Chalopin, Naville, Plard, Galiana, & Volff, 2015). Similarly, the high fraction of DNA transposons (∼10%) is in line with DNA transposon content in other fish species (Chalopin et al., 2015) but is substantially higher that what has been reported in mammals (∼3%) (Chalopin et al., 2015; Lander et al., 2001).

**Fig. 3.**
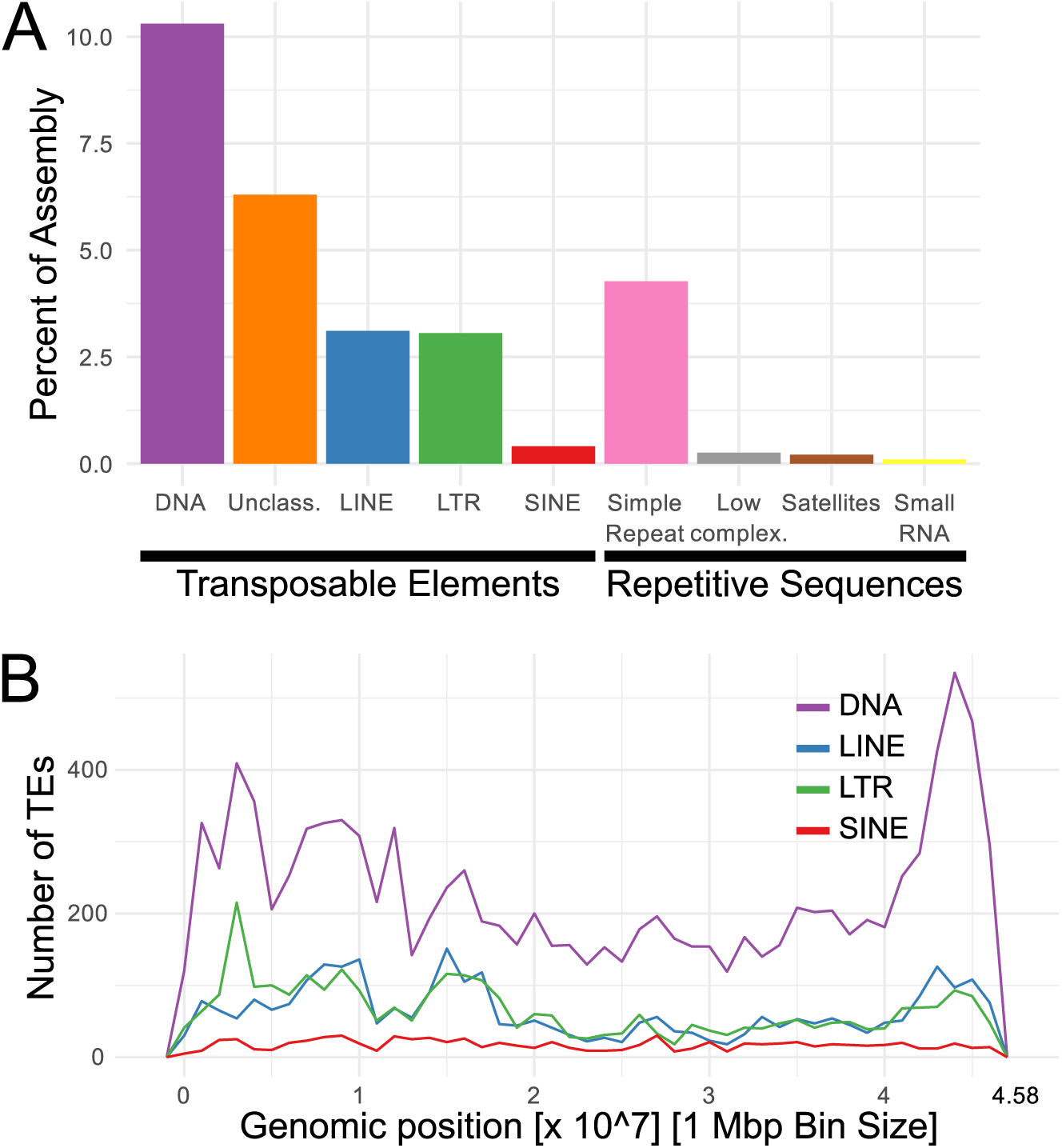
Repeat content of the orange clownfish genome assembly. **(A)** Repeat content of the whole genome as classified into transposable elements and repetitive sequences. **(B)** Spatial distribution of the four main identified classes of transposable elements on chromosome 1. Transposable element spatial distribution for chromosomes 2-24 is show in Fig S2. Detailed transposable element content is shown in Table S4.

Following the characterization of repetitive sequences in the Nemo v1 genome assembly, gene annotation was performed with the BRAKER1 pipeline, which trained the AUGUSTUS gene predictor with supplied RNA-seq data, and a successive refinement with the MAKER2 pipeline. We provided BRAKER1 with mapped RNA-seq data from 10 different tissues. This initial annotation comprised 49,881 genes with 55,273 transcripts. The gene finder models of SNAP and AUGUSTUS were refined based on the initial annotation, and MAKER2 was then used to improve the annotation using the new models and the available protein homology and RNA-seq evidence. The resulting annotation contained 26,606 genes and 35,498 transcripts, which feature a low mean AED of 0.12, indicating a very good agreement with the provided evidence. After retaining only genes with evidence support (AED of less than 1) or an annotated Pfam protein domain, the filtered annotation was comprised of 26,597 genes, corresponding to 35,478 transcripts (Table 4). This result is broadly consistent with the average number of genes (23,475) found in the 22 diploid fish species considered in this study (Table S3). Compared to the initial annotation, genes in the final annotation are 61% longer (13,049 bp) and encode mRNAs that are 80% longer (17,727 bp). The proportion of the genome that is covered by coding sequences also increased to 8.1% in the final annotation. Together with the observed reduction in the gene number by 47%, this indicates a substantial reduction of likely false positive gene annotations of short length and/or few exons. The gene density across the 24 chromosomes of our assembly varied from 23.6 genes/Mb (chromosome 21) to 36.5 genes/Mb (chromosome 14), with a genome-wide average of one gene every 29.7 Mb (Table 3). The spatial distribution of genes across all 24 chromosomes is relatively even (Fig. 1B), with regions of very low gene density presumably corresponding to centromeric regions. We observed that the longest annotated gene was APERC1_00006329 (26.5 kb), which encodes the extracellular matrix protein FRAS1, while the gene coding for the longest protein sequence was APERC1_00011517, which codes for the 18,851 amino acid protein, Titin. Functional annotation was carried out using Blast2GO and yielded annotations for 22,507 genes (85%) after aligning the protein sequences to the UniProt/Swissprot database and annotating protein domains with InterProScan.

**Table 4.** Gene annotation statistics

|  | Initial<br>BRAKER1 | Final<br>MAKER2 |
| --- | --- | --- |
| Genes | 49,881 | 26,597 |
| mRNAs | 55,273 | 35,478 |
| Exons | 391,637 | 463,688 |
| Introns | 336,364 | 428,210 |
| CDSs | 55,273 | 35,478 |
| Overlapping genes | 2,407 | 1,852 |
| Contained genes | 744 | 463 |
| Longest gene | 264,684 | 264,684 |
| Longest mRNA | 264,684 | 264,684 |
| Mean gene length | 8,097 | 13,049 |
| Mean mRNA length | 9,841 | 17,727 |
| % of genome<br>covered by genes | 44.4 | 38.2 |
| % of genome<br>covered by CDS | 7.5 | 8.1 |
| Exons per mRNA | 7 | 13 |
| Introns per mRNA | 6 | 12 |
| **BUSCO** |  |  |
| Completeness | 95.94% | 96.25% |
| Complete | 4,398 | 4,412 |
| Single-Copy | 3,588 | 3,888 |
| Duplicated | 810 | 524 |
| Fragmented | 138 | 96 |
| Missing | 48 | 76 |
| Total | 4,584 | 4,584 |

### Identification of orange clownfish-specific genes

To investigate the gene space of the orange clownfish relative to other fishes, we used OrthoFinder v1.1.4 (Emms & Kelly, 2015) to identify orthologous relationships between the protein sequences of the orange clownfish and four other fish species (Asian seabass, Nile tilapia, southern platyfish and zebrafish) from across the teleost phylogenetic tree (Betancur-R. et al., 2013). The vast majority of sequences (89%) could be assigned to one of 19,838 orthogroups, with the remainder identified as “singlets” with no clear orthologs. We observed a high degree of overlap of protein sequence sets between all five species, with 75% of all orthogroups (14,783) shared amongst all species (Fig. 4A). The proteins within these orthogroups presumably correspond to the core set of teleost genes. Of the 14,783 orthogroups with at least one sequence from each species, a subset of 8,905 orthogroups contained only a single sequence from each species. The phylogeny obtained from these single-copy orthologous gene sequences (Fig. 4B) is consistent with the known phylogenetic tree of teleost fishes (Betancur-R. et al., 2013). Interestingly, we identified a total of 4,429 sequences that are specific to the orange clownfish, 2,293 (49%) of which possess functional annotations (Fig. 4A). Future investigations will focus on the characterization of these unique genes and what roles they may play in orange clownfish phenotypic traits.

**Fig. 4.**
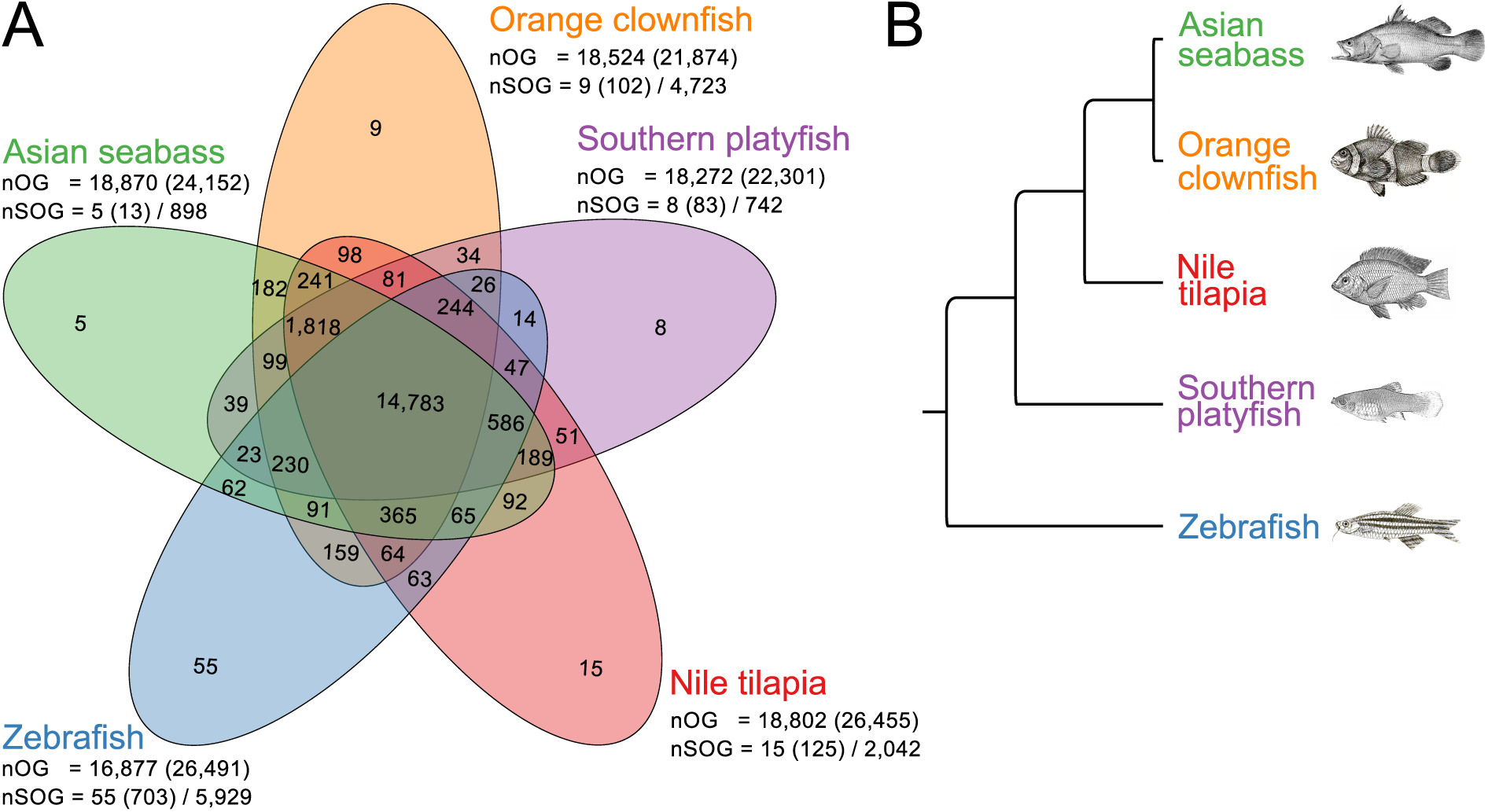
**(A)** The overlap of orthologous gene families of the orange clownfish, southern platyfish, Nile tilapia, zebrafish and Asian seabass. The total number of orthogroups (nOG) followed by the number of genes assigned to these groups is provided below the species name. The number of species-specific orthogroups (nSOG) and the respective number of genes is also indicated, followed by the number of genes not assigned to any orthogroups. **(B)** The inferred phylogenetic tree based on the ortholog groups that contain a single gene from each species.

## Conclusion

Here, we present a reference-quality genome assembly of the iconic orange clownfish, *A. percula*. We sequenced the genome to a depth of 121X with PacBio long reads and performed a primary assembly with these reads utilizing the Falcon_Unzip algorithm. The primary assembly was polished to yield an initial assembly of 903.6 Mb with a contig N50 value of 1.86 Mb. These contigs were then assembled into chromosome-sized scaffolds using Hi-C chromatin contact maps, followed by gap-filling with the PacBio reads, to produce the final reference assembly, Nemo v1. The Nemo v1 assembly is highly contiguous, with contig and scaffold N50s of 3.12 and 38.4 Mb, respectively. The use of Hi-C chromatin contact maps allowed us to scaffold 890.2 Mb (98%) of the 908.2 Mb final assembly into the 24 chromosomes of the orange clownfish. An analysis of the core set of Actinopterygii genes suggests that our assembly is nearly complete, containing 97% of the core set of highly conserved genes. The Nemo v1 assembly was annotated with 26,597 genes with an average AED score of 0.12, suggesting that most gene models are highly supported.

The high-quality Nemo v1 reference genome assembly described here will facilitate the use of this now genome-enabled model species to investigate ecological, environmental and evolutionary aspects of reef fishes. To assist the research community, we have created the Nemo Genome DB database, http://nemogenome.org/ (Fig. 5), where researchers can access, mine and visualize the genomic and transcriptomic resources of the orange clownfish.

**Fig. 5.**
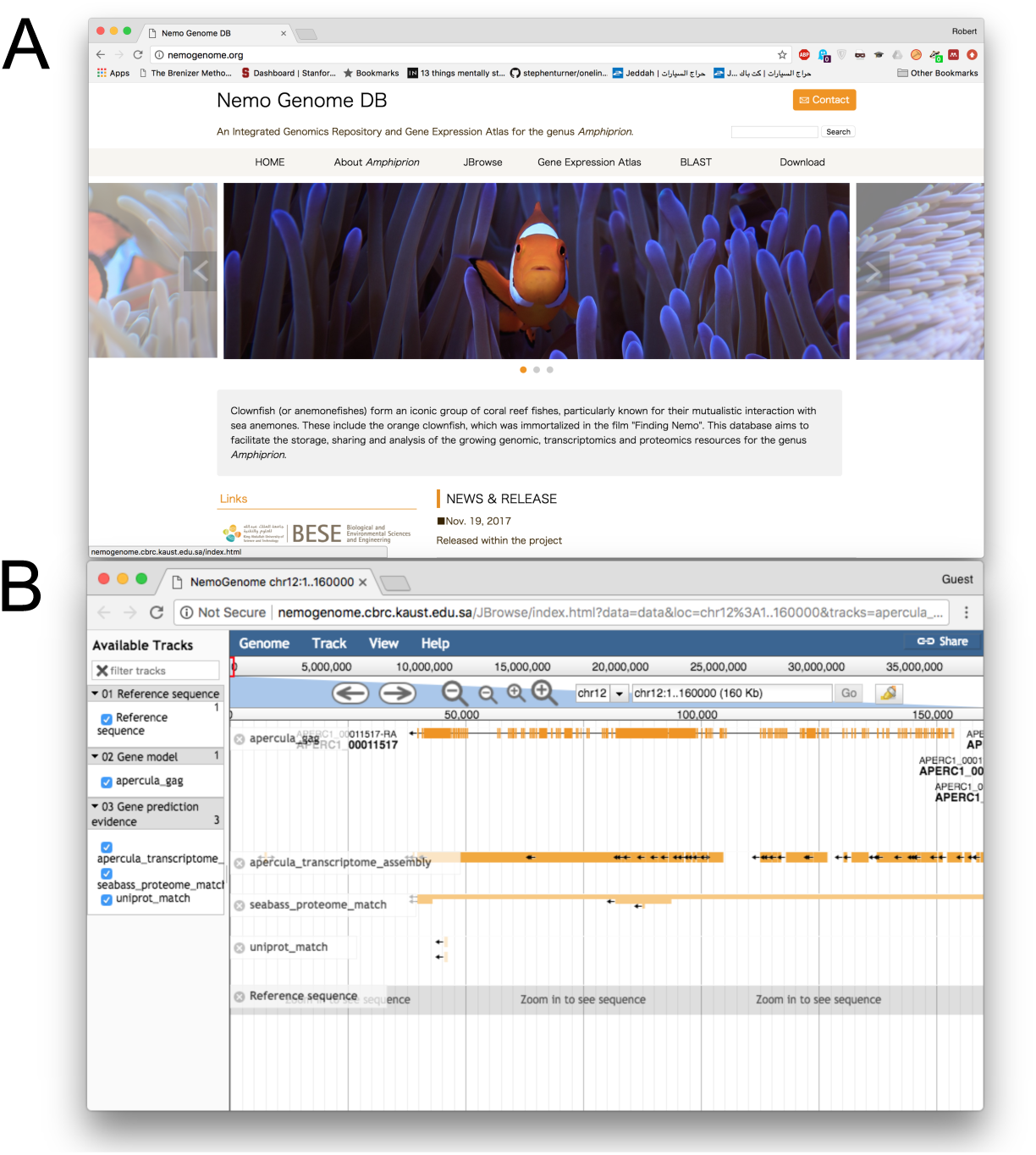
**(A)** Front page of the Nemo Genome DB database, which is a portal to access the data described in this manuscript and is accessible at www.nemogenome.org. **(B)** Genome viewer representation of the Titin gene.

## Acknowledgements

This study was supported by the Competitive Research Funds OCRF-2014-CRG3-62140408 from the King Abdullah University of Science and Technology (KAUST) to T.R., M.L.B. and P.L.M., as well as KAUST baseline support to M.L.B., M.A., T.G. and T.R. This project was completed under JCU Ethics A1233 and A1415. We thank Dr. Jennifer Donelson and staff at JCU’s MARFU facility for assistance with animal husbandry, Dr. Susanne Sprungala for DNA extraction for Illumina library preparation, KAUST BCL for the PacBio sequencing, Dr. Hicham Mansour for sequencing advice and Dr. Rita Bartossek for the PacBio library preparations. We thank Dr. Salim Bougouffa for stimulating discussions. We also acknowledge Mr. Tane Sinclair-Taylor for providing the photograph of the orange clownfish (Fig. 1A). This paper is dedicated to our good friend and colleague, Dr. Sylvain Foret.

## Author contributions

R.L. and D.J.L. designed and performed the computational analysis. R.L., T.R., C.S. and D.J.L. interpreted the results. H.O., K.M. and T.G. created the database. C.T.M. and S.F. produced sequencing libraries. R.L., D.J.L, T.R., P.L.M., M.L.B., M.A. and D.J.M. wrote the manuscript and all authors approved the final version. T.R. supervised the project.

## Data Accessibility

The assembled and annotated genome as well as the raw PacBio reads and Illumina reads are available at the Nemo Genome DB (http://nemogenome.org). Furthermore, the assembled genome will be available on GenBank as BioProject PRJNA436093 and BioSample accession SAMN08615572. All raw sequencing data described in this study will be available via the NCBI Sequencing Read Archive.

